# Arpin regulates migration persistence by interacting with both tankyrases and the Arp2/3 complex

**DOI:** 10.1101/2021.03.16.435563

**Authors:** Gleb Simanov, Irene Dang, Artem I. Fokin, Ksenia Oguievetskaia, Valérie Campanacci, Jacqueline Cherfils, Alexis M. Gautreau

## Abstract

During cell migration, protrusion of the leading edge is driven by the polymerization of Arp2/3-dependent branched actin networks. Migration persistence is negatively regulated by the Arp2/3 inhibitory protein Arpin. To better understand Arpin regulation in the cell, we looked for interacting partners and identified both Tankyrase 1 and 2 (TNKS) using a yeast two hybrid screen and co-immunoprecipitation with full-length Arpin as a bait. Arpin interacts with ankyrin repeats of TNKS through a C-terminal binding site on its acidic tail overlapping with the Arp2/3 binding site. To uncouple the interactions of Arpin with TNKS and Arp2/3, we introduced point mutations in the Arpin tail and attempted to rescue the increased persistence of the Arpin knock-out using random plasmid integration or compensating knock-in at the *ARPIN* locus. Arpin mutations impairing either Arp2/3- or TNKS-interaction were insufficient to fully abolish Arpin activity. Only the mutation that affects both interactions rendered Arpin completely inactive, suggesting the existence of two independent pathways, by which Arpin controls migration persistence. Arpin was found to dissolve liquid-liquid phase separation of TNKS upon overexpression. Together these data suggest that TNKS might be mediating the function of Arpin rather than regulating Arpin.

## 1. Introduction

Cell migration depends on various types of membrane protrusions. Most membrane protrusions are driven by cortical actin polymerization (Ridley 2011). In untransformed cells, adherent membrane protrusions driven by branched actin networks fuel cell movement at the leading edge. These branched actin networks are formed by the Arp2/3 complex that nucleates actin filaments from the side of pre-existing ones. Arp2/3 activation in membrane protrusions is under the control of the small GTPase Rac1 and, downstream, of the WAVE complex (Steffen, Koestler, and Rottner 2014; Molinie and Gautreau 2017).

Arpin was identified as an Arp2/3 inhibitory protein that antagonizes WAVE activity (Dang et al. 2013). Arpin is composed of a folded domain and of a C-terminal acidic tail protruding from this core (Fetics et al. 2016). Through its acidic tail, Arpin competes with Arp2/3 activators such as WAVE (Dang et al. 2013). Migration persistence is the result of feedback loops that sustain Rac1 activation, where Rac1 had previously induced the formation of branched actin (Krause and Gautreau 2014; Dimchev et al. 2021). By inhibiting Arp2/3, Arpin interrupts this positive feedback and decreases migration persistence, thus allowing cells to pause and change direction (Dang et al. 2013; Gorelik and Gautreau 2015).

To better understand how Arpin is regulated in the cell, we looked for Arpin interacting partners and identified TNKS, as major Arpin partners. TNKS are pleiotropic regulators of many cell functions through the binding, modification and down-regulation of a myriad of proteins. However, TNKS did not appear to regulate Arpin and its Arp2/3 inhibitory function, but rather to serve as another Arpin effector in the regulation of migration persistence.

## 2. Results

### 2.1 Arpin binds to Tankyrase 1 and 2

To identify proteins that bind to Arpin, we first immunoprecipitated Arpin from a stable 293 cell line expressing a Protein C (PC) tagged version of Arpin through its epitope tag. Arpin was efficiently immunoprecipitated from the lysate prepared from cells stably expressing tagged Arpin, but not from a control cell line transfected with the empty plasmid. Silver staining was used to identify potential interacting partners (Fig.1A). In a single step immunoprecipitation, many bands were detected, including immunoglobulin light and heavy chains. However, two specific bands between the 120 and 150 kDa markers were clearly detected when Arpin was immunoprecipitated, but not in the control lane. These two proteins were identified by LC-MS/MS as Tankyrase1 (TNKS1, 142 kDa) and Tankyrase2 (TNKS2, 127 kDa). With the expectation to identify additional potential partners of the Arpin protein, we performed a yeast two hybrid screen of a library containing random primed human placenta cDNAs, with full-length Arpin as a bait. More than 10^8^ clones were analyzed by yeast mating. Out of the 187 clones selected, 177 corresponded to either TNKS1 or TNKS2 (Fig.1B). The remaining 10 clones were comparatively of low confidence, as they corresponded to out-of-frame fusions or to DNA sequences that were not annotated as protein encoding genes. These two approaches thus point at TNKS as major Arpin partners.

**Figure 1.**
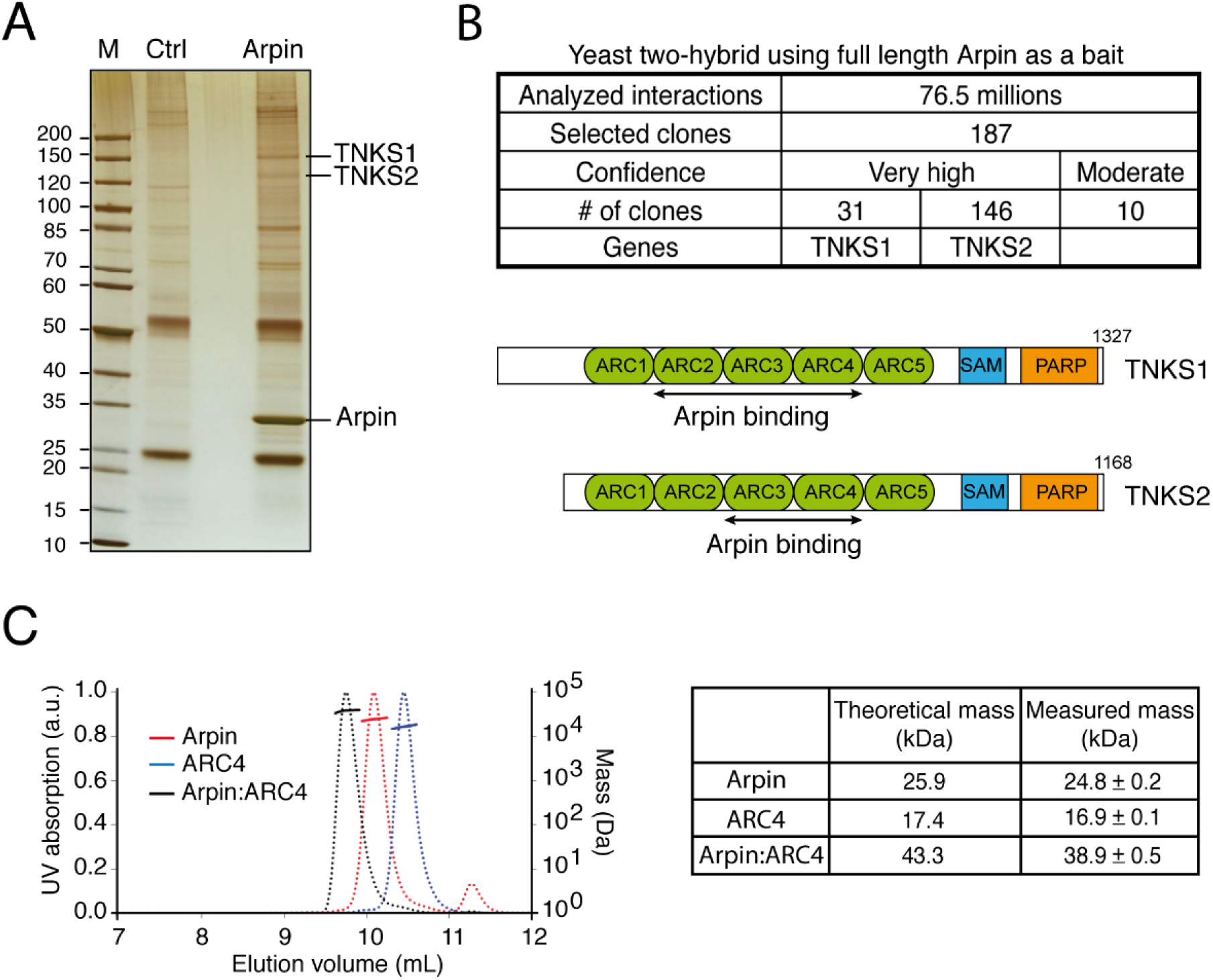
Tankyrases are major Arpin interacting partners. (**A**) A 293 stable cell line expressing tagged Arpin was used to immunoprecipitate Arpin and associated proteins (silver staining). Partner proteins were identified by mass spectrometry. (**B**) Full-length Arpin was used as a bait in a yeast two-hybrid screen. The retrieved clones of TNKS mapped to regions indicated with black arrows within the region containing Ankyrin Repeat Clusters (ARCs). (**C**) The molecular species obtained by mixing purified Arpin and purified ARC4 of TNKS were analyzed by SEC-MALS.

Both TNKS are composed of three regions. From N- to C-terminus, these proteins contain ankyrin repeats organized into Ankyrin Repeat Clusters (ARCs), a Sterile Alpha Motif (SAM), which mediates oligomerization, and a C-terminal Poly ADP Ribosyl Polymerase (PARP) catalytic domain (Hsiao and Smith 2008). Since all yeast two-hybrid clones interacting with full-length Arpin mapped to the N-terminal region composed of ARCs, we produced and purified full-length Arpin and the ARC4 of TNKS2, which has been previously crystallized (Guettler et al. 2011). When the two proteins were mixed, a new molecular species was detected by Size Exclusion Chromatography – Multi Angle Light Scattering (SEC-MALS). As expected, this species displays a mass corresponding to a 1:1 complex (Fig.1C).

### 2.2 Arpin levels do not appear to be regulated by TNKS

TNKS are pleiotropic regulators of various cellular functions, including telomere maintenance, mitosis regulation, Wnt signaling, insulin-dependent glucose uptake, the Hippo-YAP pathway (Smith 1998; P. Chang, Coughlin, and Mitchison 2005; W. Chang, Dynek, and Smith 2005; Riffell, Lord, and Ashworth 2012; Kim, Dudognon, and Smith 2012; S.-M. A. Huang et al. 2009; Wang et al. 2015). These multiple TNKS functions usually require binding to substrates through ARCs and poly ADP ribosylation, also called PARylation, through the catalytic PARP domain (Eisemann et al. 2016). Usually, but not always, the fate of PARylated proteins is to be ubiquitinated by the E3 ubiquitin ligase RNF146, which recognizes poly-ADP ribose chain through its WWE domain, and then to be degraded by proteasomes (Y. Zhang et al. 2011).

TNKS turnover fast, because they PARylate themselves. To investigate whether Arpin turns over through a similar mechanism, we blocked TNKS catalytic activity with the XAV939 inhibitor (S.-M. A. Huang et al. 2009). As expected, this treatment resulted in increased levels of both TNKS and of their substrate Axin1. Levels of Arpin was, however, not modified by the XAV939 treatment (Fig.2A). This observation is in line with proteomics analyses of TNKS function: Levels of Arpin (referred in these large-scale studies as C15ORF38) were also found to be unchanged in TNKS double knock-out versus control 293T cells (Bhardwaj et al. 2017) and the Arpin-TNKS interaction was unaffected when TNKS PARylation activity was blocked by XAV939 or not (X. Li et al. 2017). We nonetheless attempted to detect the potential PARylation of Arpin. To this end, we used the WWE domain of E3 ligase RNF146 as a recognition module to pull down PARylated proteins (Y. Zhang et al. 2011). In the WWE pull-down, TNKS1, but not Arpin, was retrieved (Fig.2B).

**Figure 2.**
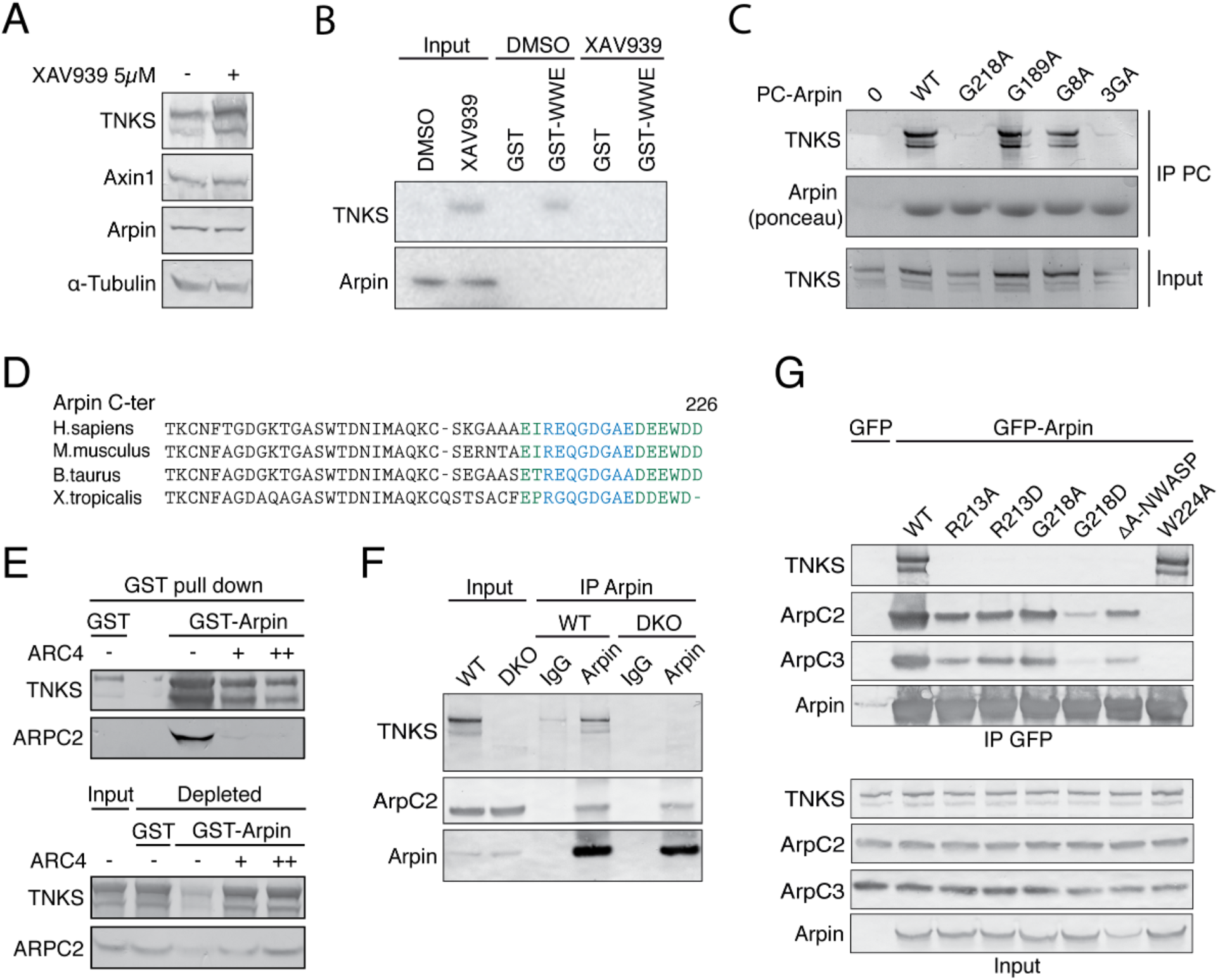
The Arpin-TNKS interaction does not appear to regulate Arpin levels and requires a C-terminal consensus motif for TNKS binding. This consensus motif overlaps with Arp2/3 interaction site but TNKS do not modulate the Arpin-Arp2/3 interaction in cells. (**A**) The Arpin-TNKS interaction does not appear to regulate Arpin levels. 293T cells were treated with the TNKS inhibitor XAV939, or vehicle, for 24h, and the lysates were analyzed by Western blots. (**B**) MEF lysates were subjected to GST pulldown using the WWE domain of E3 ligase RNF146, which recognizes PARylated proteins. XAV939 (1µM) was used to block TNKS catalytic activity. (**C**) Mapping of the TNKS binding on Arpin. PC tagged Arpin WT, G8A, G189A, G218A and triple mutant G8A-G189A-G218A were transiently expressed in 293T cells. Anti-PC agarose beads were used to immunoprecipitate tagged Arpin. (**D**) TNKS (blue) and Arp2/3 (green) binding sites overlap in the C-terminus acidic tail of Arpin. (**E**) TNKS binding competes Arp2/3 binding in vitro. MEF lysates were incubated with purified GST- or GST-Arpin immobilized on glutathione beads. Increasing concentrations of purified ARC4 was added to the lysate as indicated. GST beads, input lysate and depleted lysates were analyzed by Western blots. (**F**) Lysates from WT and TNKS double KO 293T cells were analyzed by Arpin immunoprecipitations or non-immune IgG as a control. (**G**) GFP-Arpin WT, R213A, R213D, G218A, G218D, ΔA-NWASP and W224A were transiently expressed in 293T cells and immunoprecipitated.

Together these experiments indicate that Arpin is a direct TNKS partner, which is unlikely to be subjected to PARylation-mediated degradation. The situation of Arpin contrasts with the majority of TNKS binding partners, but is similar to several previously described TNKS binding partners that are not PARylated, such as Mcl-1L, GDP Mannose 4,6 Dehydratase, CD2AP, SSSCA1 (Bae, Donigian, and Hsueh 2003; Bisht et al. 2012; Kuusela et al. 2016; Perdreau-Dahl et al. 2020).

### 2.3. Arpin binds to TNKS via its C-terminal acidic tail

ARCs recognize a consensus motif, the octapeptide RXXXXGXX, defined through the screening of a peptide library (Guettler et al. 2011). Arpin contains three putative TNKS binding sites, which were examined by substituting the required G residue by A at positions 8, 189 and 218. We expressed PC tagged Arpin mutant forms in 293T cells and noticed that the G218 residue is the only critical one for the ability of Arpin to associate with TNKS (Fig.2C). This TNKS binding motif is located in the acidic tail of Arpin and overlaps with the previously described Arp2/3 interaction site (Fig.2D) (Dang et al. 2013). We therefore investigated a possible competition between Arp2/3 complex and TNKS binding. GST pull-down with Arpin in lysates from mouse embryonic fibroblasts (MEF) retrieved the Arp2/3 complex and both TNKS (Fig.2E). Adding an excess of purified ARC4 did not only displace TNKS, but also the Arp2/3 complex, in line with overlapping binding sites. In order to understand whether TNKS can influence Arpin-Arp2/3 interaction in cells, we immunoprecipitated endogenous Arpin from 293T wild type and TNKS double knock-out (KO) cells (Bhardwaj et al. 2017). The same amount of Arp2/3 complex co-precipitated with Arpin whether TNKS were present or not (Fig.2F). Thus, in cells, the Arpin-TNKS interaction does not appear to modulate the Arpin-Arp2/3 interaction. Since TNKS do not regulate the levels of Arpin, nor its Arp2/3 inhibitory function, we then investigated what might be the role of the Arpin-TNKS interaction.

We attempted to uncouple TNKS and Arp2/3 binding using mutations of the Arpin acidic tail. To prevent TNKS binding, we replaced R213 and G218 by A or D residues. Alanine substitutions are most classical to impair binding sites, but introducing aspartate is a way to increase negative charges of the Arpin tail, a requisite for Arp2/3 binding (Pollard 2007). On the opposite, to impair Arp2/3 binding, we substituted the conserved C-terminal tryptophan of Arpin by alanine (W224A). We expressed these mutant forms of Arpin in 293T cells and immunoprecipitated them to analyze their binding partners (Fig.2G). As expected, TNKS interaction was undetectable when R213 or G218 of Arpin were mutated. The Arp2/3 interaction was below the detection limit with the W224A substitution. It was also surprisingly affected in the G218D substitution, even though this mutation increased the overall acidity of the tail. G218D thus impaired both TNKS and Arp2/3 binding.

### 2.4 Arpin controls the ability of TNKS to form biomolecular condensates

TNKS can oligomerize via their SAM motif. The polymeric state reinforces the PARP activity of TNKS and is required for Wnt signaling (Mariotti et al. 2016; Riccio et al. 2016). Furthermore, when overexpressed, TNKS form cytosolic aggregates (De Rycker and Price 2004; Mariotti et al. 2016; Riccio et al. 2016; X. Li et al. 2017). We found that the aggregates formed by GFP-TNKS2 in transiently transfected MCF10A cells (Fig.3A) were dynamic, since they can fuse (Fig.3B), and recover fast fluorescence after photobleaching (Fig.3C). TNKS2 aggregates recovered up to 70 % of their intensity in 100 s. These properties suggest that GFP-TNKS2 aggregates undergo liquid-liquid phase separation and form so-called biomolecular condensates (Alberti, Gladfelter, and Mittag 2019). Such condensates are often controlled by multivalent interactions. TNKS display 5 ARCs for each protomer of a multimer. The TNKS partners that display several binding sites, such as Axin (Morrone et al. 2012), can bring together several multimeric TNKS units and may thereby promote a liquid-liquid phase transition.

**Figure 3.**
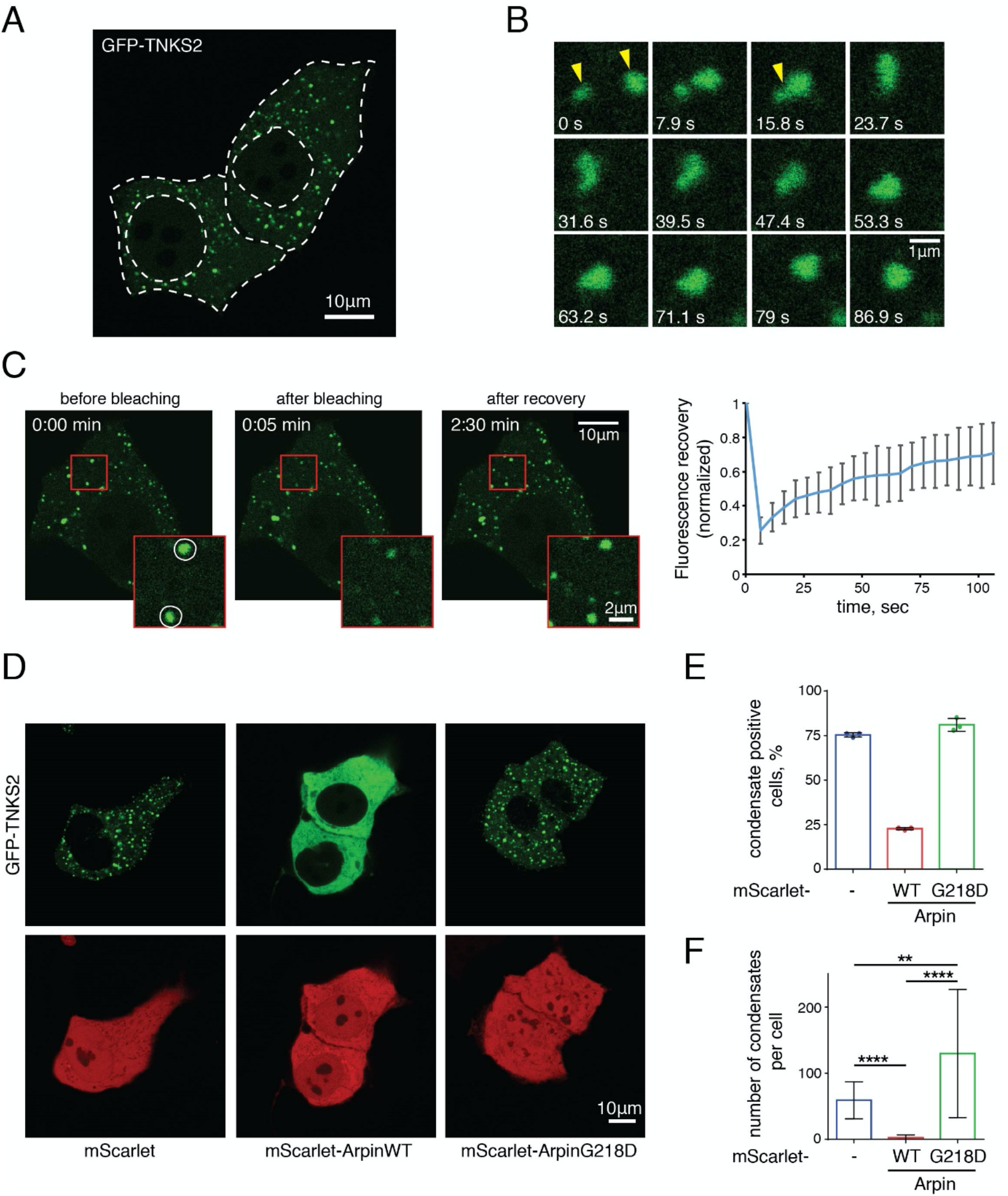
Arpin regulates the ability of TNKS to form biomolecular condensates. (**A**) GFP-TNKS2 was transiently expressed in MCF10A cells. White dashed lines delimit cell membranes and nuclei. (**B**) Fusion of two GFP-TNKS2 aggregates (yellow arrows) was imaged in the time-lapse series. (**C**) GFP-TNKS2 aggregates were analyzed by FRAP (bleached areas are indicated by white circles in the magnified image boxed in red). (**D**) Wild type or the G218D Arpin fused to mScarlet were co-expressed with GFP-TNKS2 in MCF10A cells. (**E**) Quantification of the proportion of transfected cells that display TNKS2 condensates (3 biological replicates). (**F**) Quantification of the number TNKS2 condensates per cell (Kruskal-Wallis test, ** *p* < 0.01, **** *p* < 0.0001).

We thus examined whether Arpin, which displays a single binding site, might regulate the formation of biomolecular condensates by TNKS. As a negative control, we used the G218D mutant form, which displays reduced Arp2/3 binding, in addition to its impairment of TNKS interaction. When we co-expressed Arpin with GFP-TNKS2, wild type Arpin, but not G218D Arpin, prevented the formation of TNKS condensates (Fig.3D). This behavior is different from the one of endogenous or overexpressed Axin, which was reported to co-localize with TNKS condensates (Mariotti et al. 2016; Riccio et al. 2016). Co-expression of wild type Arpin decreased the fraction of aggregate-positive cells (Fig.3E) and the number of condensates per cell (Fig.3F). Since these results depend on TNKS overexpression, we also examined whether Arpin would exert such a function at the endogenous level of TNKS expression. In MCF10A cells, TNKS are diffuse in the cytoplasm in WT and *ARPIN* KO cells (Fig.S1). Upon XAV939 treatment, TNKS condensate. No difference in TNKS condensation was observed in WT and *ARPIN* KO cells. Since the ability to form biomolecular condensates is thought to correspond to increased catalytic activity, we also examined levels of TNKS and of their substrate Axin1 and PTEN. However, these levels were unchanged in *ARPIN* KO cells compared to parental cells (Fig.S2). In conclusion, Arpin has a striking role in preventing TNKS from forming biomolecular condensates when both components are overexpressed, but the implication of this result at the endogenous level of expression is not clear.

### 2.5 The Arpin-TNKS interaction participates to the regulation of cell migration

To examine the role of the Arpin-TNKS interaction, we used the MCF10A *ARPIN* KO cell line (Molinie et al. 2019) to isolate clones stably re-expressing WT Arpin or derivatives. We first focused on two Arpin mutations, G218A and W224A, that impair TNKS and Arp2/3 binding respectively (Fig.2G). Exogenous Flag tagged Arpins were moderately overexpressed compared to the endogenous Arpin (Fig.4A). A major role of the Arp2/3 pathway in cell migration is to mediate migration persistence through positive feedback and the Arp2/3 inhibitory protein Arpin antagonizes this role (Dang et al. 2013; Krause and Gautreau 2014). We recorded random migration of single cells using these cell lines. As previously reported (Molinie et al. 2019), *ARPIN* KO cells exhibited higher migration persistence compared to parental MCF10A cells. Expression of wild type Arpin fully rescued the *ARPIN* KO phenotype (Fig.4B). The W224A mutant form, which is impaired in its interaction with Arp2/3 partially rescued the phenotype, but almost as efficiently as wild-type. The G218A mutant form, that still binds to Arp2/3 but not to TNKS, also partially rescued the phenotype, but less efficiently than W224A. All migration parameters extracted from cell trajectories are displayed in figure S4 for reference, but the only parameter that is regulated by Arpin in all cell systems is migration persistence, not speed, nor mean square displacement (Dang et al. 2013; Krause and Gautreau 2014; Molinie et al. 2019). These results suggested that the Arpin-TNKS interaction can regulate migration persistence, but we sought to confirm them in cell clones, where Arpin is not overexpressed.

**Figure 4.**
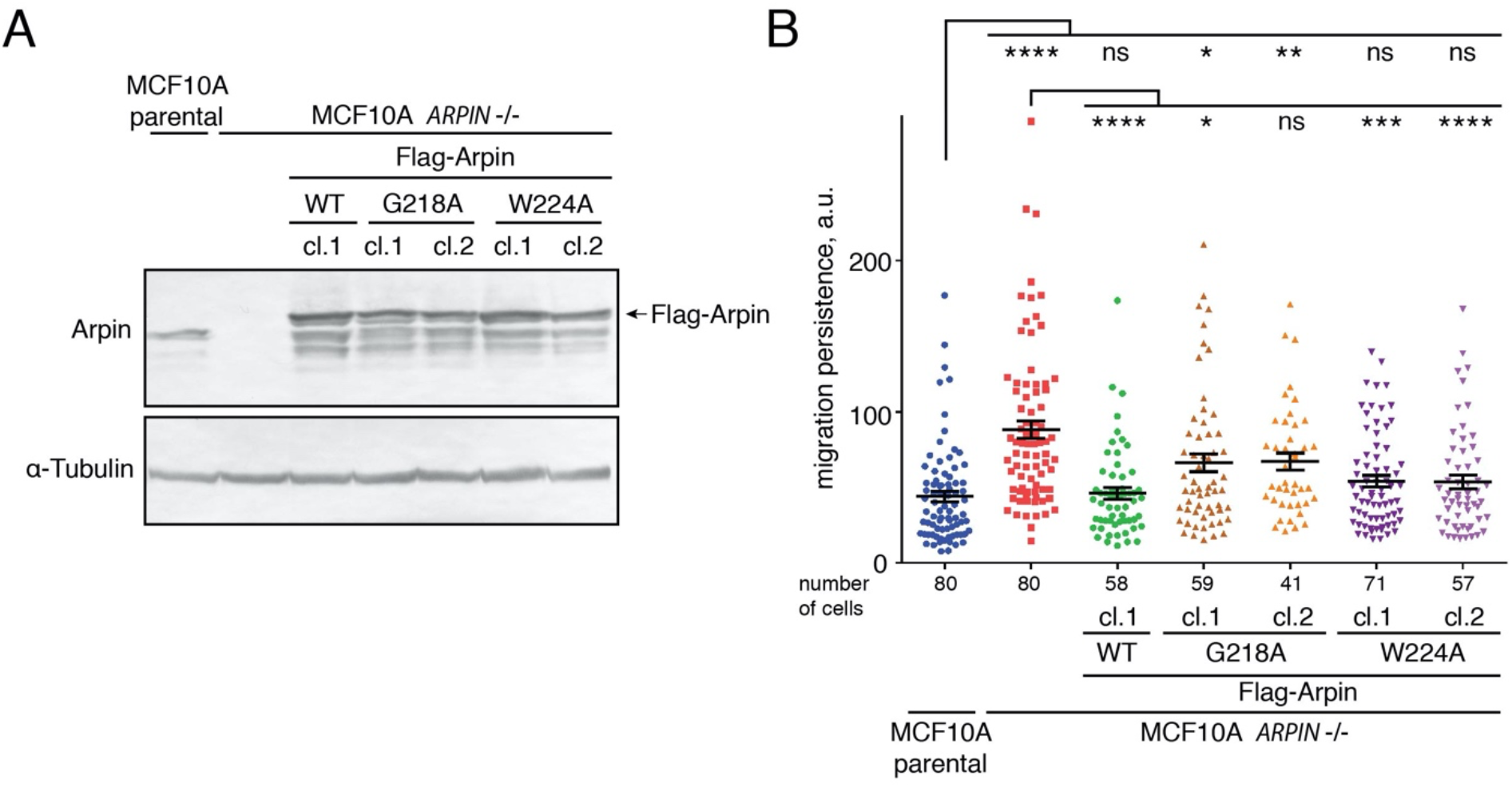
The Arpin-TNKS interaction is important for the control of cell migration persistence. (**A**) Stable clones expressing Flag-Arpin WT, Flag-Arpin G218A and Flag-Arpin W224A after plasmid random integration were generated from the *ARPIN* KO MCF10A cell line. Endogenous and exogenous Arpins were revealed by Western blot using Arpin antibodies. (**B**) Random migration of single cells was tracked for 7 h and migration persistence was extracted from trajectories (Kruskal-Wallis test, * *p* < 0.05, ** *p* < 0.01, **** *p* < 0.0001, number of cells ranges from 41 to 80).

For this purpose, we designed a GFP-Arpin Knock-In (KI) strategy. Briefly, we introduced two double strand breaks (DSBs) to excise the exons encoding the *ARPIN* open reading frame from *ARPIN* KO MCF10A cells and provided a donor plasmid encoding GFP-Arpin WT, G218A, G218D, W224A for Homology-Directed Repair (HDR; Fig.5A and Methods section). GFP-Arpin expression was confirmed by Western blot in stable clones isolated upon puromycin selection (Fig.5B). Indeed, transgene expression was overall at the level of the endogenous, even though differences could still be observed between constructs and clones. The various GFP-Arpin forms appeared mostly diffuse in the cell, but low levels of expression made live cell imaging difficult. We performed immunofluorescence staining of fixed cells using Arpin and secondary antibodies coupled to organic fluorophores to enhance the signal. All GFP-Arpins were indeed similarly cytosolic and nuclear, like the endogenous Arpin (Fig.S4).

We then performed the single cell migration assay with our KI clones. WT Arpin fully rescued the *ARPIN* KO phenotype (Fig.5C). In the KI system also, we observed rescue with W224A Arpin. Rescue with W224A Arpin was more or less efficient depending on the clone and this variation was not an effect of expression levels. The G218A Arpin provided a partial rescue, that failed to reach significance, indicating that this mutant that abolished TNKS binding is more severely affected than W224A in the KI as well as in the overexpression system. Only the G218D Arpin that impaired the interaction with both Arp2/3 and TNKS was completely unable to rescue the *ARPIN* KO phenotype. All migration parameters extracted from cell trajectories are displayed in figure S5 for reference. These results suggest that the interactions of Arpin with Arp2/3 and TNKS represent two pathways that both regulate cell migration.

**Figure 5.**
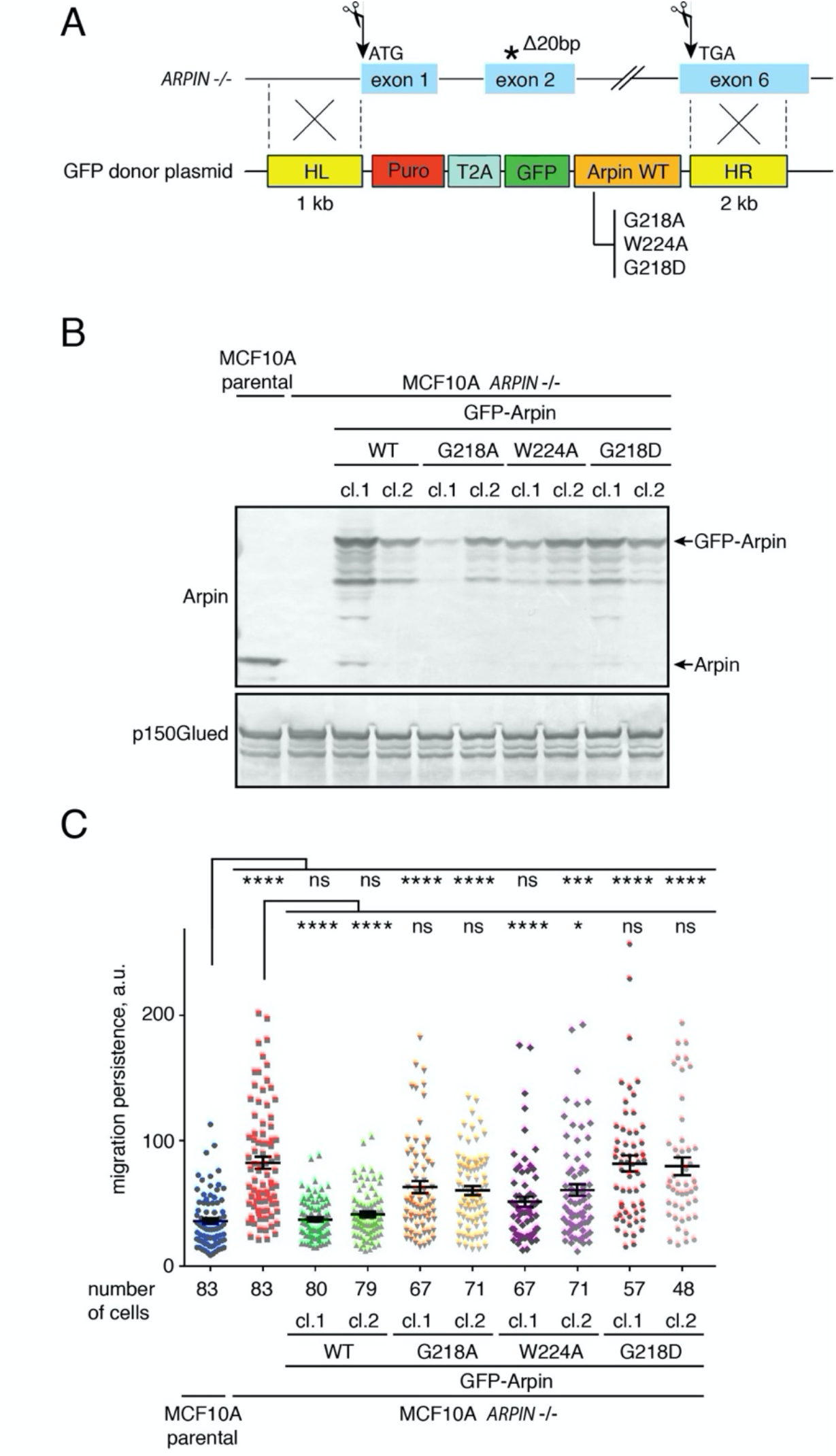
The Arpin-TNKS interaction controls cell migration persistence. (**A**) Scheme of the knock-in strategy used to rescue *ARPIN* KO cells. Two gRNAs allowing the excision of the whole Arpin locus were designed. The premature stop codon that appears due to a 20 bp deletion is indicated by a *. Homology-directed repair was used to integrate selectable donor cassettes. The cassettes contain a single Open Reading Frame encoding the puromycin resistance gene, the viral self-cleaving T2A peptide, GFP and Arpin WT, G218A, W224A or G218D. (**B**) Expression of GFP-Arpin WT and mutant forms in selected cell lines was assessed by Western blot using Arpin antibodies. (**C**) Migration persistence was extracted from single cells migrating randomly (Kruskal-Wallis test, * *p* < 0.05, *** *p* < 0.001, **** *p* < 0.0001, number of cells ranges from 47 to 83).

## 3. Discussion

Here we report that TNKS are major Arpin partners in the cell. TNKS bind to the exposed C-terminal tail that protrudes from a folded core domain (Fetics et al. 2016). The C-terminal tail carries the previously reported Arp2/3 inhibitory binding site (Dang et al. 2013). The TNKS binding site overlaps the one of Arp2/3 on Arpin tail and we found that one ARC of TNKS can displace the Arp2/3 bound to the tail of Arpin in vitro. However, this competition does not appear to take place in the cell, since the amount of Arp2/3 bound to Arpin does not increase in TNKS double KO cells. Arpin is thought to bind Arp2/3 at the lamellipodial edge upon Rac1 signaling. The results obtained here rather suggest that Arpin bind to TNKS in a diffuse manner in the cytosol or the nucleus. In the cell, the lack of competition of Arp2/3 and TNKS for Arpin binding might be due to the fact that these two partners do not bind Arpin in the same location.

Our analysis of point mutations of the Arpin tail suggests that Arp2/3 and TNKS are both important for the regulation of migration persistence. Indeed, point mutations that specifically impair one or the other of Arpin partner display only a partial loss of activity, even in a clean KI context associated with endogenous levels of expression. In contrast, the G218D mutation that significantly impairs binding to both Arp2/3 and TNKS is clearly loss of function. Previously, we had reported that the deletion of the whole C-terminal tail fully inactivated Arpin (Dang et al. 2013). This is consistent with our current results, but it can no longer be interpreted as the sole lack of Arp2/3 binding. TNKS binding to Arpin participates to the regulation of migration persistence independently of Arp2/3 binding.

TNKS were previously implicated in the regulation of cell migration. Since TNKS are overall overexpressed in several cancer types and are promising targets to block in particular the Wnt pathway, pharmacological inhibition of TNKS or their siRNA-mediated depletion was tested and shown in numerous studies to decrease migration and invasion of cancer cell lines (Bao et al. 2012; Kang et al. 2012; Tian et al. 2014; Lupo et al. 2016; C. Li et al. 2018; Ha et al. 2018; Yang et al. 2019; J. Huang et al. 2020). Given the plethora of TNKS partners, the mechanisms at play may not be the same in all cell systems. One TNKS partner, TNKS1BP1, which is PARylated, negatively regulates cancer cell invasion by interacting with the capping protein and decreasing actin filament dynamics (Ohishi et al. 2017). TNKS1BP1 is downregulated in pancreatic cancer.

Here we report that TNKS aggregates fulfill properties of biomolecular condensates and that Arpin dissolves these condensates upon overexpression. Biomolecular condensates correspond to liquid-liquid phase separation due to multimeric proteins and multimeric ligands (P. Li et al. 2012). TNKS possess multiple ARCs and oligomerize through their SAM motif (Mariotti et al. 2016; Riccio et al. 2016). The presence of multivalent ligands induce TNKS condensation (Diamante et al. 2021). On the contrary, Arpin is a monomeric protein with a single TNKS binding site that fits very well the consensus motif defined by peptide display library (Guettler et al. 2011). Overexpressed Arpin is thus likely to saturate functional ARCs of TNKS, resulting in the displacement of endogenous multivalent ligands and hence dissolution of TNKS condensates. However, at endogenous levels of expression, we neither detected a role of Arpin in TNKS condensation, nor in the efficiency with which TNKS regulate their substrates. So it is still unclear at this point how Arpin controls TNKS and how TNKS control migration persistence.

Biomolecular condensation might play a role in tumor cells, where TNKS are overexpressed. However, it should be stressed that Arpin is, on the contrary, down-regulated in tumors compared to normal adjacent tissue (Lomakina et al. 2016; Liu et al. 2016; T. Li et al. 2017; S.-R. Zhang et al. 2019). Since Arpin and TNKS levels vary in opposite directions in cancers, the here reported interaction between Arpin and TNKS might be more important in the regulation of cell migration in untransformed cells than in tumor cells. In untransformed cells, such as MCF10A cells, Arpin appears to control migration persistence through a two-pronged mechanism, involving the independent binding of Arp2/3 and TNKS to the same binding site of Arpin.

## 4. Materials and Methods

### 4.1 Plasmids, gRNAs and transfection

For expression in 293 Flp-In cells, human Arpin ORF was cloned in pcDNA5 FRT His PC TEV Blue between FseI and AscI sites. 293 Flp-In stable cell line expressing PC Arpin was obtained as previously described (Derivery and Gautreau 2010). For the yeast two-hybrid screen, full length human Arpin was cloned in pB27 in fusion with the LexA DNA binding domain. A random primed cDNA library from human placenta was screened by Hybrigenics using a mating protocol and 2 mM 3-aminotriazole to reduce background. Arpin G8A, G189A, G218A, triple G8A-G189A-G218A, R213A, R213D, G218D, W224A mutants were obtained in the pcDNA5 His PC TEV Arpin plasmid using QuikChange Lightning Site-Directed Mutagenesis Kit (Agilent). ORFs encoding Arpin WT and mutant forms were subcloned between FseI and AscI sites into custom-made pcDNAm FRT PC GFP, MXS PGK ZeoM bGHpA EF1Flag mScarlet Blue2 SV40pA, and MXS EF1Flag Blue2 SV40pA PGK Blasti bGHpA plasmids.

293T cells were transfected using Calcium Phosphate or Lipofectamine 3000 (Thermo Fisher Scientific). The 293T *ARPIN* KO cell line (clone #44) was generated with CRISPR/Cas9 system, as previously described for MCF10A cells (Molinie et al. 2019). Stable MCF10A cells expressing Flag-Arpin WT, Flag-Arpin G218A and Flag-Arpin W224A were obtained in MCF10A *ARPIN* KO cells (Molinie et al. 2019) by transfecting with custom-made MXS plasmids described above using Lipofectamine 3000 (Thermo Fisher Scientific). Cells were selected with 10 µg/ml of Blasticidin (InvivoGen). Individual clones were picked with cloning rings and Flag-Arpin expression was checked by Western blot. MCF10A Arpin KI cell lines were generated with CRISPR/Cas9 system. Following targeting sequences were used: 5’-TCCCGACCGCCCGGGCACCC-3’ targets before ATG codon in exon1, 5’-GATTTCTCTAGGATGACTGA-3’ targets after Stop codon in exon6 of Arpin. These sequences were flanked by BbsI restriction site. Corresponding oligonucleotides were annealed and cloned in the pX330 plasmid expressing human SpCas9 protein (Addgene #42230). The donor plasmids were constructed as follows. Sequences were amplified by PCR with Phusion polymerase (Thermo Fisher Scientific): Arpin homology arm right (HR) flanking Cas9 targeted site was amplified from genomic DNA extracted from wild type MCF10A cells (NucleoSpin tissue extraction kit, Macherey-Nagel), Puro-T2A were amplified from the custom-made plasmid MXS Puro bGHpA using primers containing T2A sequence. Amplified sequences were checked by Sanger sequencing. Arpin homology arm left (HL) was synthesized by Eurofins. The donor cassette was constructed by assembling HL, Puro-T2A, GFP-Blue2 and HR by MXS-Chaining (Sladitschek and Neveu 2015). Full length ORFs encoding Arpin WT, G218A, G218D, W224A were then subcloned in the constructed donor plasmid between FseI and AscI sites. MCF10A *ARPIN* KO cells were transfected with the Cas9- and gRNA-containing pX330 plasmid and the donor plasmids described above using Lipofectamine 3000 (Thermo Fisher Scientific). Cells were selected with 0.5 µg/ml of puromycin (InvivoGen). Single clones were picked with cloning rings, expanded and analyzed by Western blot.

### 4.2 Cell culture and drugs

MCF10A cells were maintained in DMEM/F12 medium (Thermo Fisher Scientific) supplemented with 5% horse serum (Sigma), 100 ng/ml cholera toxin (Sigma), 20 ng/ml epidermal growth factor (Sigma), 0.01 mg/ml insulin (Sigma), 500 ng/ml hydrocortisone (Sigma) and 100 U/ml penicillin/streptomycin (Thermo Fisher Scientific). 293T and HeLa cells were maintained in DMEM medium (Thermo Fisher Scientific) supplemented with 10% fetal bovine serum (Thermo Fisher Scientific) and 100 U/ml penicillin/streptomycin (Thermo Fisher Scientific). Cells were incubated at 37 °C in 5% CO2. All cells and stable clones were routinely tested for mycoplasma and found to be negative. TNKS inhibitor XAV939 was from Sigma. 293T parental and TNKS double KO cell lines were kindly provided by Dr. S. Smith, Skirball Institute, New York School of Medicine.

### 4.3 Immunoprecipitation and GST pull-down

Stable 293 cells expressing His-PC-Arpin or the empty plasmid as a control (Fig.1) were lysed in (50 mM Hepes pH7.7, 150 mM NaCl, 1mM CaCl2, 1% NP40, 0.5% Na Deoxycholate, 0.1% SDS 1% supplemented with protease inhibitor cocktail, Roche). Clarified lysates were incubated with 10 µl of HPC4 coupled beads (Sigma) for 3 h at 4°C. After 5 washes in the same buffer, beads were analyzed by SDS-PAGE. For TNKS identification, tryptic peptides were analyzed by NanoLC-MS/MS analyses using a LTQ Orbitrap Velos mass spectrometer (Thermo Scientific) coupled to the EASY nLC II high performance liquid chromatography system (Proxeon, Thermo Scientific). Peptide separation was performed on a reverse phase C18 column (Nikkyo Technos). NanoLC-MS/MS experiments were conducted in a Data Dependent acquisition method by selecting the 20 most intense precursors for CID fragmentation and analysis in the LTQ. Data were processed with the Proteome Discoverer 1.3 software and protein identification was performed using the Swissprot database and MASCOT search engine (Matrix science).

For PC immunoprecipitation of PC-Arpin WT or mutants (Fig.2), 293T cell lysates were lysed in (50 mM KCl, 10 mM Hepes pH 7.7, 1 mM MgCl_2_, 1 mM EGTA, 1% Triton TX100 supplemented with protease inhibitor cocktail, Roche). 20µl of HPC4 beads were supplied with 1mM Ca^2+^, protease inhibitor cocktail (Roche) and 2µM ADP HDP PARG inhibitor (MerckMillipore). Beads were incubated with extracts for 1 h at 4°C, washed 5 times in the same buffer and analyzed by Western blot. For GFP immunoprecipitation (Fig.2), 293T cells transiently transfected with GFP-Arpin were lysed in (50 mM KCl, 10 mM Hepes pH 7.7, 1 mM MgCl_2_, 1 mM EGTA, 1% Triton TX100 supplemented with protease inhibitor cocktail, Roche). Extracts were incubated with anti-GFP agarose beads (GFP-trap, Chromotek) for 2h at 4°C, washed 5 times in the same buffer and analyzed by Western blot.

For immunoprecipitation of endogenous Arpin, 293T cells were lysed in (50 mM KCl, 10 mM Hepes pH 7.7, 1 mM MgCl_2_, 1 mM EGTA, 1% Triton TX100 supplemented with protease inhibitor cocktail, Roche). Clarified extracts were incubated for 2 h with agarose beads previously coupled to 10 µg of non-immune rabbit IgG or 10 µg of affinity purified Arpin antibodies (according to the manufacturer protocol, AminoLink Coupling Resin, Thermo Fisher Scientific). Beads were incubated with extracts for 2 h at 4°C, washed 5 times in the same buffer and analyzed by Western blot. HeLa cell pellets were lysed in (50 mM Hepes pH7.7, 150 mM NaCl, 1mM CaCl2, 1% NP40, 0.5% Na Deoxycholate, 0.1% SDS 1% supplemented with protease inhibitor cocktail, Roche). 20 µg of GST fusion protein and 20 µl of Glutathione Sepharose 4B Beads (GE Healthcare) were incubated with 1 ml of HeLa cell extract for 2 h at 4°C. When indicated, purified ARC4 protein is added into the mixture to compete the interaction. Beads were washed 5 times in the same buffer and analyzed by Western blot.

### 4.4 SEC-MALS

For SEC-MALS, purified proteins were separated in a 15 ml KW-803 column (Shodex) run on a Shimadzu HPLC system. MALS, QELS and RI measurements were achieved with a MiniDawn Treos, a WyattQELS and an Optilab T-rEX (all from Wyatt technology), respectively. Mass calculations were performed with the ASTRA VI software (Wyatt Technology) using a dn/dc value of 0.183 mL.g^-1^.

### 4.5 Protein purification and analysis of Arpin expression

To produce recombinant proteins in *E.coli*, ARC4 from TNKS2 was cloned into pQE30 (Qiagen). Arpin and the WWE domain of RNF146 (amino-acids 100-175) were produced as GST fusion proteins from a modified pGEX vector containing a TEV protease cleavage site after the GST moiety. For analysis of Arpin expression MCF10A cells were lysed in RIPA buffer (50mM Hepes, pH 7,5, 150mM NaCl, 1% NP-40, 0.5% DOC, 0,1% SDS, 1mM CaCl2) supplemented with the EDTA free protease inhibitor cocktail (Roche), the lysates were clarified and analyzed by Western blot.

### 4.6 Western blots and Antibodies

SDS-PAGE was performed using NuPAGE 4-12% Bis-Tris gels (Life Technologies). For Western blots, proteins were transferred using the iBlot system (Life Technologies) and developed using HRP-coupled antibodies, Supersignal kit (Pierce) and a LAS-3000 imager (Fujifilm) or AP-coupled antibodies and NBT/BCIP as substrates (Promega). Rabbit polyclonal antibodies obtained and purified using full-length Arpin were previously described (Dang et al. 2013). The following commercial antibodies were used: TNKS1/2 pAb (H-350, Santa Cruz Biotechnology), ArpC2 pAb (Millipore), ArpC3 pAb (Sigma-Aldrich), α-Tubulin mAb (Sigma), Axin1 mAb (C76H11 Cell Signaling Technology), PTEN pAb (Cell Signaling Technology, #9552).

### 4.7 Immunofluorescence

For immunofluorescence MCF10A cells were seeded on glass coverslips coated with 20 µg/mL bovine fibronectin (Sigma) for 1h at 37 °C in PBS. Then cells were fixed with 4% paraformaldehyde, permeabilized in 0.5 % Triton X-100 and blocked in 0.1 % Triton X-100, 2% bovin serum albumin (BSA, Sigma) in PBS for 1h at RT. Coverslips were incubated with indicated primary antibodies for 1h at RT, then with anti-rabbit Alexa Fluor 488 (Invitrogen). Imaging was performed on the Axio Observer microscope (Zeiss) with the 63x/1.4 oil objective.

### 4.8 Live confocal imaging

GFP-TNKS2 condensates were imaged using the confocal laser scanning microscope (TCS SP8, Leica) equipped with the inverted frame (Leica), the high NA oil immersion objective (HC PL APO 63×/1.40, Leica) and the white light laser (WLL, Leica). The acquisition was performed using LASX software. To capture condensates fusion events GFP-TNKS2 expressing cells were imaged every 0.79 s for 158 s. For the FRAP experiment single GFP-TNKS2 condensates were bleached during 1.5 s and imaged every 5 s for 105 s. The aggregates intensity was manually normalized to the background along the acquisition, the recovery curve presents the mean intensity and standard deviation (SD) (n = 30 condensates from 16 cells). To measure the fraction of cells with detectable condensates (n > 1) MCF10A cells co-expressing GFP-TNKS2 and mScarlet, mScarlet-Arpin WT or G218D mutant were imaged (data from 3 independent experiments with at least 115 cells analyzed for each condition is represented). Total quantity of GFP-TNKS2 aggregates per cell was measured in 3 independent experiments. Images were analyzed in ImageJ as follows, first, the MaxEntropy threshold was applied, then the number of condensates was counted with Analyse Particles function (size > 0.15µm; measures from 60 cells were pooled).

### 4.9 Live imaging and Analysis of cell migration and statistics

MCF10A cells were seeded onto glass bottomed µ-Slide (Ibidi) coated with 20 µg/ml of fibronectin (Sigma). Imaging was performed on the Axio Observer microscope (Zeiss) equipped with the Plan-Apochromat 10x/0.25 air objective, the Hamamatsu camera C10600 OrcaR2 and the Pecon Zeiss incubator XL multi S1 RED LS (Heating Unit XL S, Temp module, CO2 module, Heating Insert PS and CO2 cover). Pictures were taken every 10 min for 24 h. Single cell trajectories were obtained by tracking cells with Image J and analyzed using the DiPer software (Gorelik and Gautreau 2014) to obtain migration parameters: directional autocorrelation, mean square displacement, average cell speed, single cell trajectories plotted at origin. Data from two (Flag-Arpin cell lines) or three (GFP-Arpin knock-in cell lines) independent experiments were pooled for the analysis and plotted. Results are expressed as means and standard errors of the mean (s.e.m). The average cell speed was analyzed using GraphPad software with the Kruskal-Wallis test. For migration persistence statistical analysis was performed using R. Persistence, measured as movement autocorrelation over time is fit for each cell by an exponential decay with plateau (as described in (Polesskaya et al. 2020))

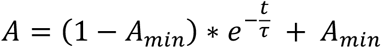

where *A* is the autocorrelation, *t* the time interval, *A*_*min*_ the plateau and *τ* the time constant of decay. The plateau value *A*_*min*_ is set to zero for cell lines in vitro as they do not display overall directional movement. The time constant *τ* of exponential fits were then plotted and compared using Kruskal-Wallis test. Four levels of statistical significance were distinguished: * *p* < 0.05, ** *p* < 0.01, *** *p* < 0.001, **** *p* < 0.0001.

## Author Contributions

Conceptualization and methodology: G.S., A.I.F. and A.M.G.; Investigation, validation, formal analysis and visualization: G.S., I.D., K.O. and V.C. Writing of the original draft: G.S. Writing, review and editing: G.S. and A.M.G. Project administration, supervision, resources and funding acquisition: A.M.G. All authors have read and agreed to the submit the manuscript.

## Funding

This research was funded by grants from the Agence Nationale de la Recherche (ANR ANR-15-CE13-0016-01) and from Institut National du Cancer (INCA_6521).

## Acknowledgments

We thank Dr. Susan Smith for a kind gift of TNKS double KO cell line, Véronique Henriot, Anaïs Pitarch, Angelina Chemeris and Sai P. Visweshwaran for technical support, Marc Lavielle and Chuang Yu for their help with statistical tools. This work was supported by French Infrastructure for Integrated Structural Biology (FRISBI) ANR-10-INSB-05-01. We thank Manuela Argentini and Christophe Velours from the I2BC facilities in Gif-sur-Yvette for mass spectrometry analyses (SICaPS) and for SEC-MALS, respectively. We thank the Polytechnique Bioimaging Facility for assistance with live imaging on their equipment partly supported by Région Ile-de-France (interDIM) and Agence Nationale de la Recherche (ANR-11-EQPX-0029 Morphoscope2, ANR-10-INBS-04 France BioImaging).

## Conflicts of Interest

The authors declare no conflict of interest.

## Notes

### Competing Interest Statement

The authors have declared no competing interest.

