## Supplementary data for "Arpin regulates migration persistence by interacting with both tankyrases and the Arp2/3 complex"

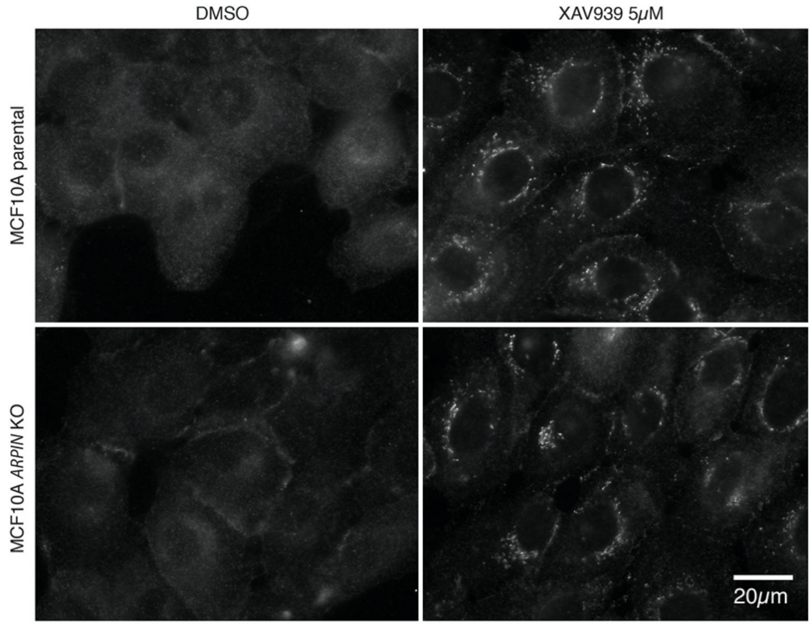


**Figure S1.** Immunofluorescence staining of TNKS in MCF10A WT and *ARPIN* KO cells treated with XAV939 at 5 µM for 9 h or with DMSO. TNKS are mostly cytosolic in both cases, but appear to condensate upon XAV939 treatment.

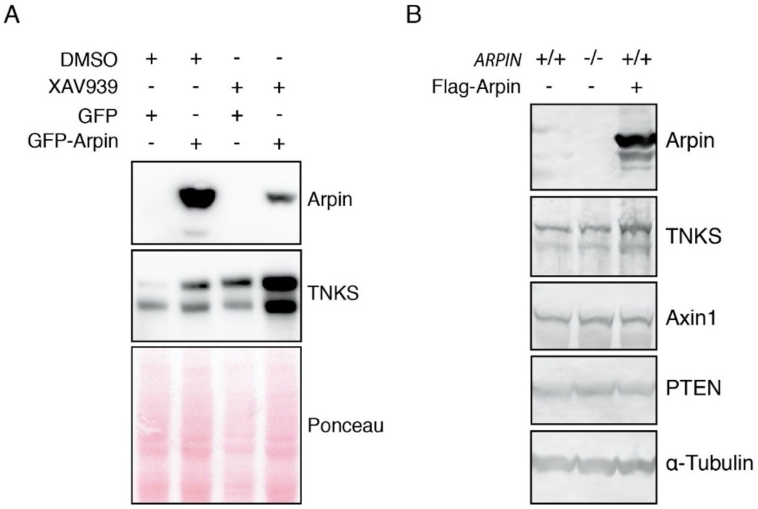


**Figure S2.** Arpin overexpression increases TNKS protein levels. (**A**) 293T cells were transiently transfected with GFP or GFP-Arpin and lysed. Where indicated, XAV939 1µM was added for 24 h (**B**) 293T WT or *ARPIN* KO cells were transiently transfected with indicated constructs and lysed. Protein levels were revealed by Western blot with corresponding antibodies.

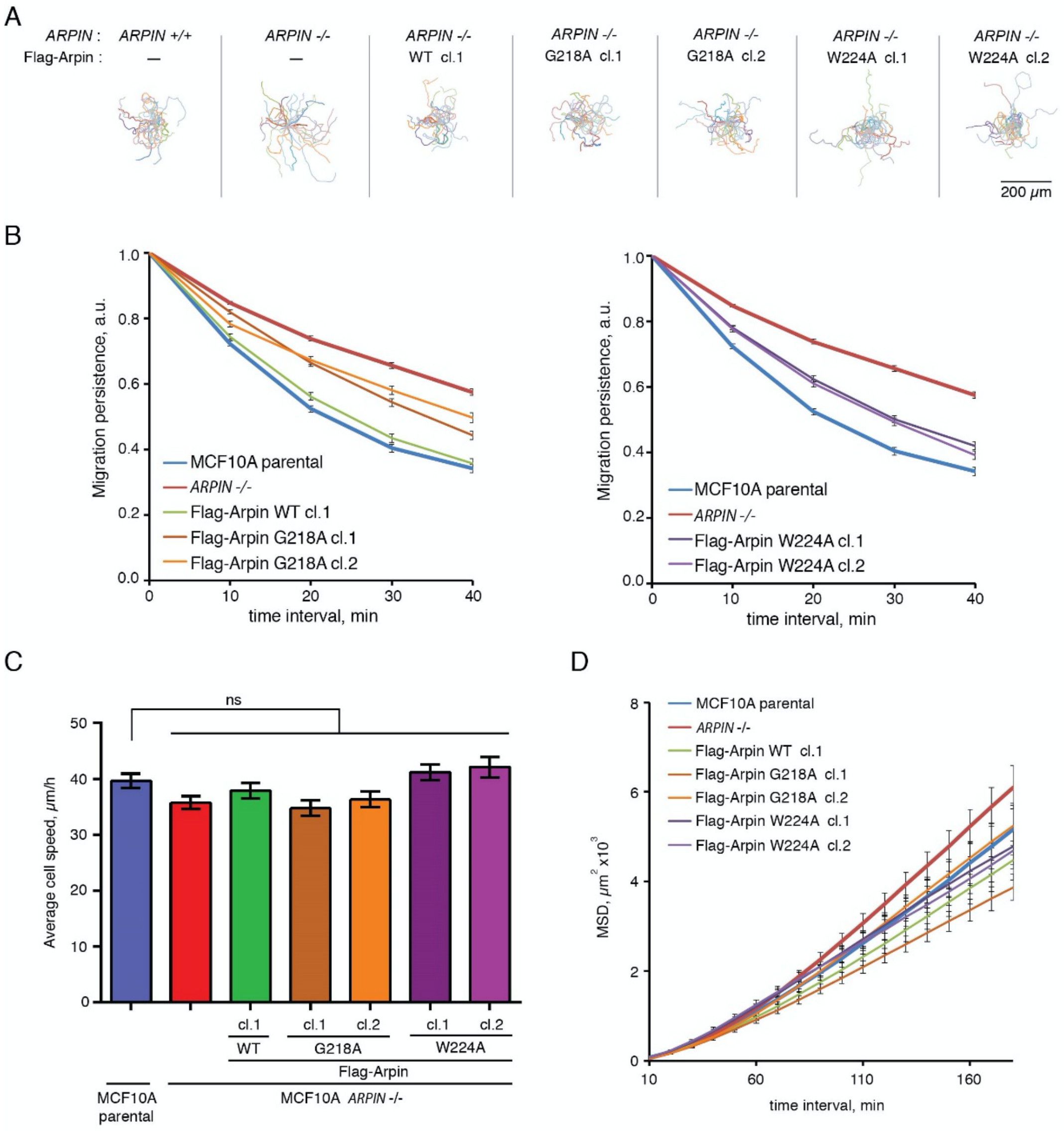


**Figure S3.** Single cell migration assay of stable MCF10A clones expressing WT, G218A and W224A Flag-Arpin. (**A**) Single cell trajectories (25 cells tracked for 7 h displayed. (**B**) Migration persistence displayed here for the whole cell population as an autocorrelation curve (Gorelik and Gautreau 2014). (**C**) Average cell speed (**D**) Mean square displacement (MSD) is represented as its mean value and standard error of the mean (number of cells ranges from 41 to 80).

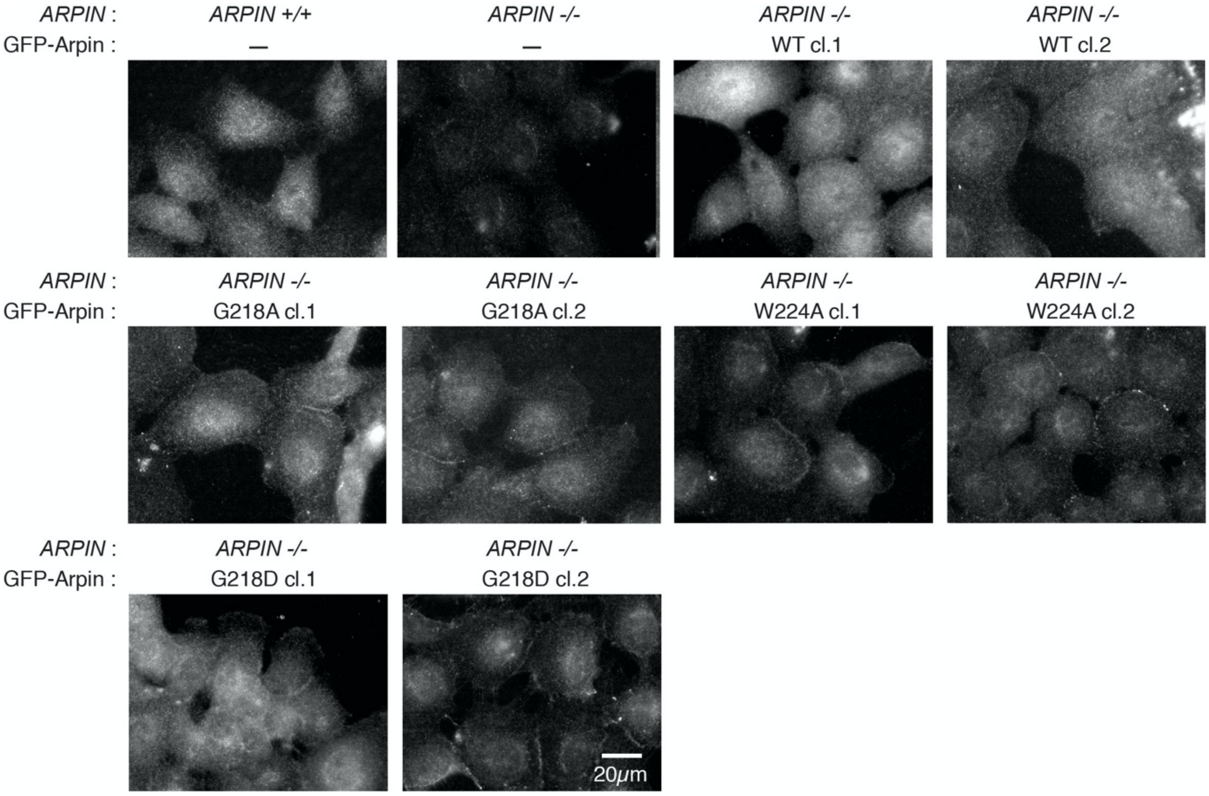


**Figure S4.** Immunofluorescence of Arpin in MCF10A parental cells, ARPIN knock-out and knock-in clones. All Arpin forms were localized in the cytosol and nucleus with an occasional peripheral staining, which does not systematically correspond either to cell junctions or to free edges.

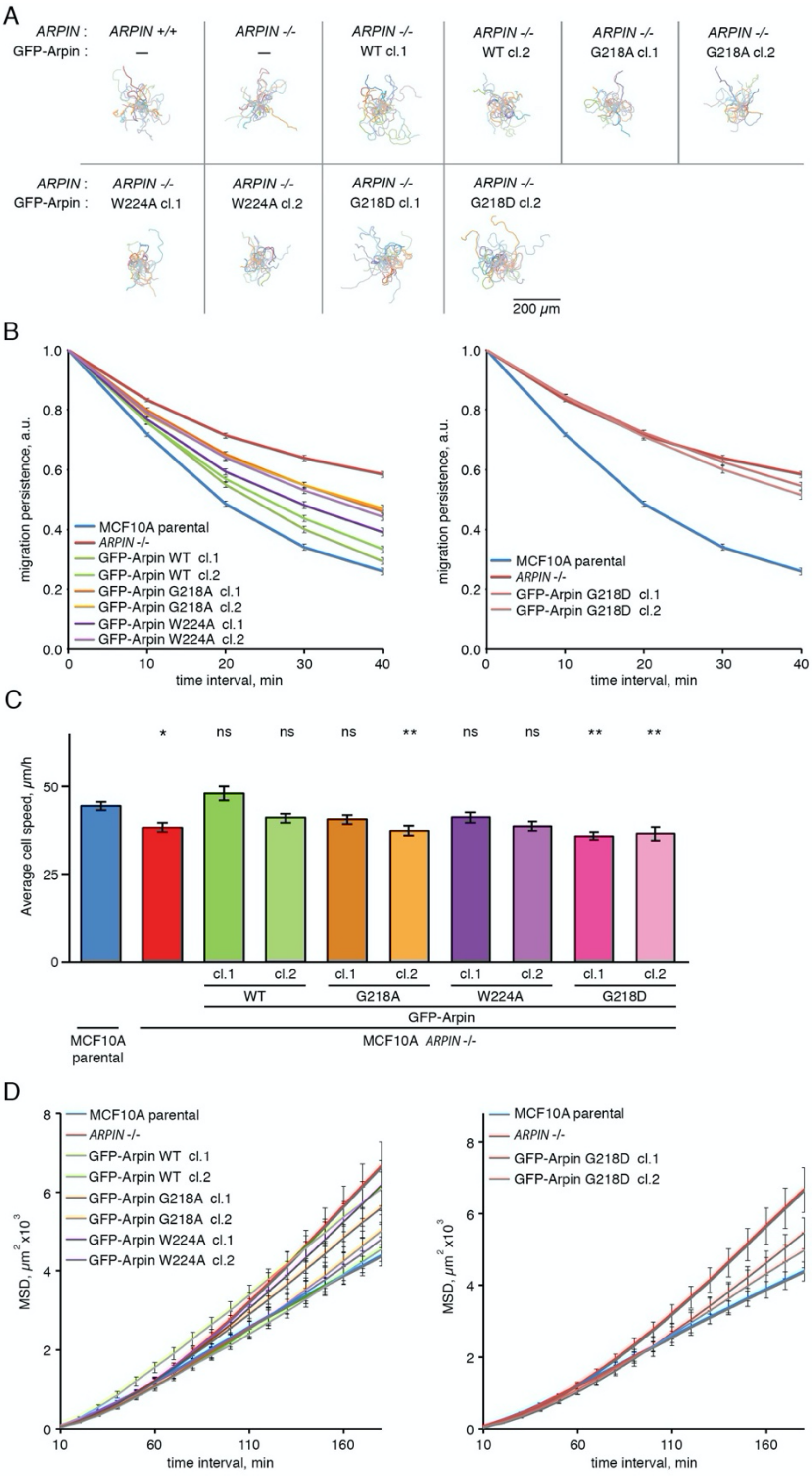


**Figure S5**. Single cell migration assay of *ARPIN* knock-in MCF10A clones. (**A**) Single cell trajectories (25 cells tracked for 7 h displayed). (**B**) Migration persistence displayed here for the whole cell population as an autocorrelation curve (Gorelik and Gautreau 2014). (**C**) Average cell speed (Kruskal-Wallis comparison with control MCF10A cells, * $p < 0.05$, ** $p < 0.01$, *** $p < 0.001$). (**D**) Mean square displacement (MSD) is represented as its mean value and standard error of the mean (number of cells ranges from 47 to 83).

**Abbreviations**

| | |
|---|---|
| ARC | Ankyrin Repeat Cluster |
| DSB | Double Strand Break |
| FRAP | Fluorescence Recovery After Photobleaching |
| KI | Knock-in |
| KO | Knock-out |
| HDR | Homology-Directed Repair |
| MEF | Murine Embryonic Fibroblast |
| PARP | Poly ADP Ribose Polymerase |
| PC | Protein C |
| SAM | Sterile Alpha Motif |
| SEC-MALS | Size Exclusion Chromatography - Multi Angle Light Scattering |
| TNKS | Tankyrases |
| WT | Wild type |
